# Innovations in double digest restriction-site associated DNA sequencing (ddRAD-Seq) method for more efficient SNP identification

**DOI:** 10.1101/2022.09.06.506835

**Authors:** Zenaida V. Magbanua, Chuan-Yu Hsu, Olga Pechanova, Mark Arick, Corrinne E. Grover, Daniel G. Peterson

## Abstract

We present an improved ddRAD-Seq protocol for identifying single nucleotide polymorphisms (SNPs). It utilizes optimally sized restriction enzyme digestion fragments, quick acting ligases that are neutral with the restriction enzyme buffer eliminating buffer exchange steps, and adapters designed to be compatible with Illumina index primers. Library amplification and barcoding are completed in one PCR step, and magnetic beads are used to purify the genomic fragments from the ligation and library generation steps. Our protocol increases the efficiency and decreases the time to complete a ddRAD-Seq experiment. To demonstrate its utility, we compared SNPs from our protocol with those from whole genome resequencing data from *Gossypium herbaceum* and *Gossypium arboreum*. Principal component analysis demonstrated that the variability of the combined data was explained by the genotype (PC1) and methodology applied (PC2). Phylogenetic analysis showed that the SNPs from our method clustered with SNPs from the resequencing data of the corresponding genotype. Sequence alignments illustrated that for homozygous loci, more than 90% of the SNPs from the resequencing data were discovered by our method. Our analyses suggest that our ddRAD-Seq method is reliable in identifying SNPs suitable for phylogenetic and association genetic studies while reducing cost and time over known methods.

## 1. Introduction

Genotyping by sequencing methods comprise a suite of techniques designed to resequence entire genomes (Eck *et al*., 2009; Cheon *et al*., 2018), multiple targeted loci, or specific genomic fractions, all with the objective of identifying single nucleotide polymorphisms (SNPs) among samples. While whole genome resequencing gives the greatest genomic coverage, it can easily become costly and analytically challenging for polyploid and/or large genome species, both of which describe many agricultural crops. Therefore, there exists interest in reliable methods to reduce genome representation while still leveraging the throughput and coverage of next generation sequencing (NGS) technologies. The first of these reduced representation techniques involved modifications of traditional gel-based methods such as restriction fragment length polymorphism (RFLP) (Keim *et al*., 1990; Schüller *et al*., 1992; Young, 1992; Ota *et al*., 2007; Brar *et al*., 2008) and amplified fragment length polymorphism (AFLP) (Liu *et al*., 1999; Mueller and Wolfenbarger, 1999; Campbell and Bernatchez, 2004; Roden, Dutton and Morin, 2009), both of which use restriction enzyme digestion to select for homologous genomic DNA fractions. In a foundational study, Baird *et al*. (2008) demonstrated a NGS adaption of these methods by digesting the genomic DNA of threespine stickleback and *Neurospora crassa* with a restriction enzyme and subsequently generating Illumina libraries from the fragments by attaching barcoded adapters and amplifying the selected fragments via PCR. The resulting sequences were termed “restriction-site associated DNA (RAD) tags”. After this initial introduction, variations on RAD tag generation were developed. Elshire *et al*. (2011) chose a methylation-sensitive restriction enzyme to generate the representative loci, which minimized the inclusion of repetitive regions, while Etter *et al*. (2011) added a second ligation step to a divergent Y-adapter (i.e., two DNA strands of adapter are complementary on only one end, forming a Y). This adapter was designed such that only fragments that were ligated to the first adapter get amplified, thereby increasing the efficiency. Subsequently, Peterson *et al*. (2012) introduced the double digest restriction-site associated DNA (ddRAD) method, which uses two restriction enzymes – a frequent and a rare cutter – and a precise size selection step prior to library amplification to further reduce representation of the genome in library construction. This method, (*i*.*e*., ddRAD-Seq) generally follows the same protocol as the single restriction enzyme methods vis-à-vis digestion, additions of adapters, amplification, and sequencing, and may include similar modifications to RAD-seq, such as using a divergent-Y in the second adapter (DaCosta and Sorenson, 2014) or using a single divergent-Y adapter and attaching the barcodes during PCR (Peterson *et al*., 2014). Wang *et al*. (2017) validated ddRAD-Seq against the SNP chip method and found more than 99% consistency.

In this study, we describe a streamlined method for ddRAD-Seq that builds upon existing work to improve both efficiency and cost-effectiveness. We use this pipeline to generate new data for multiple accessions of two cotton species, *G. herbaceum* and *G. arboreum*, and compare the results with existing whole-genome resequencing data for the same accessions. Principal component analysis (PCA) on SNPs derived from the combined data separates the samples first by genotype and subsequently by method, i.e., ddRAD-Seq data adequately represents the genome. Phylogenetic analysis of the data places the same sample from both methods into distinct clades on the tree, also suggesting high-fidelity between the ddRAD-Seq data and the whole-genome resequencing data. Importantly, comparison between the ddRAD-Seq data and the whole-genome resequencing reveals that while 67% of found SNPs were identical, over 90% of homozygous SNPs matched.

## 2. Materials and Methods

A schematic representation of the method is provided in Figure 1 (A and B).

**Figure 1.**
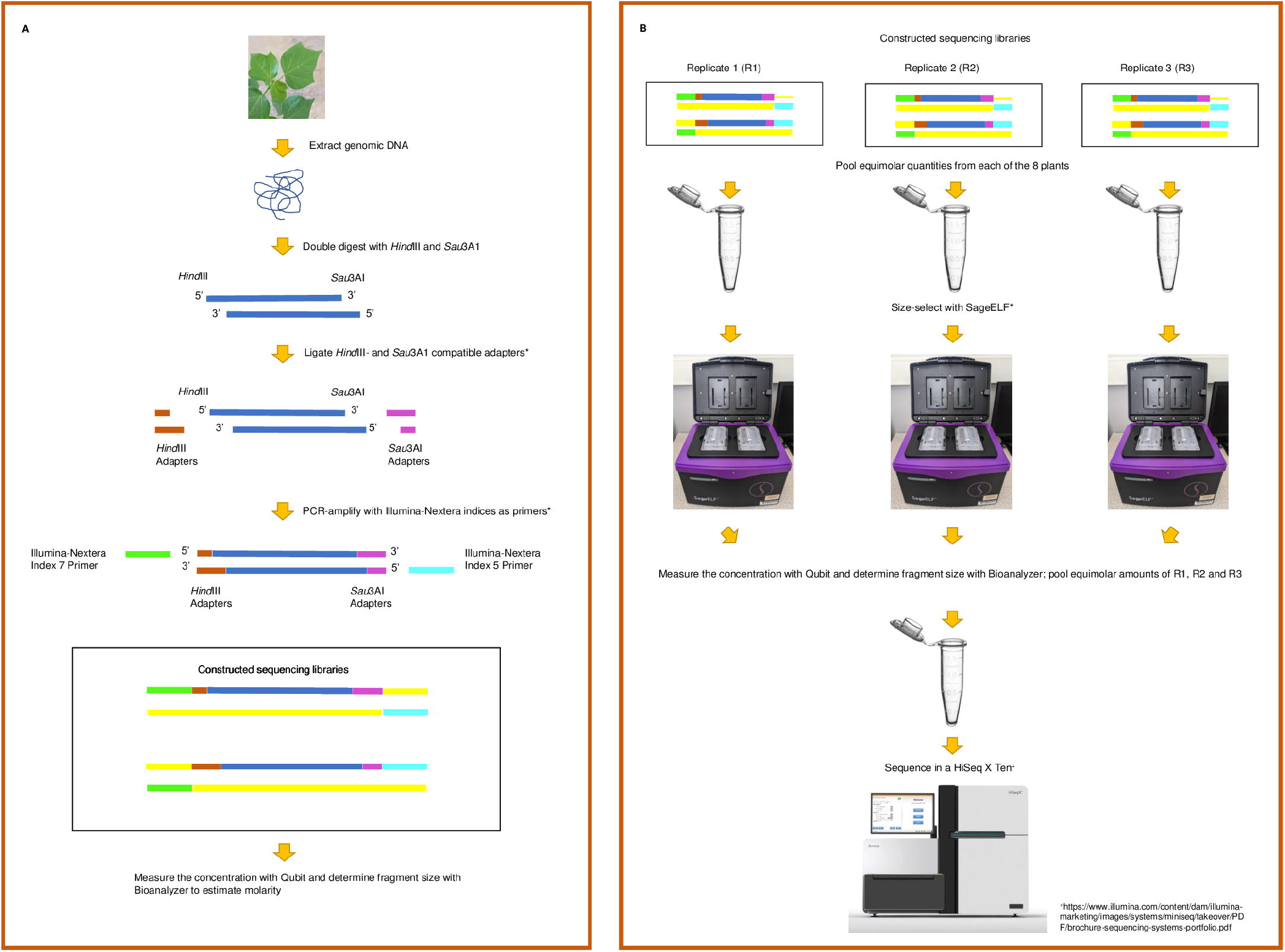
Flowchart of the methodology. (A) Shows process from genomic DNA extraction to construction of the sequencing libraries. (B) Illustrates how each replicate was treated and the point where they were pooled for sequencing. Steps with asterisks were followed with bead-purification.

### 2.1 Electronically guided digestion selection (egads)

When using Illumina technology-based ddRAD-Seq, the restriction enzyme combination should produce DNA fragments between 150 and 700 bp because this fragment range consistently gives us high quality sequence scores (Figure 1A). To facilitate selection of suitable enzyme combinations, we developed a script we refer to as “egads” which is short for electronically guided digestion selection. For a genome sequence of choice, egads predicts the likely size distribution of digestion products for a list of enzyme pairs. The script is available at https://github.com/IGBB/egads.

### 2.2 Extraction and characterization of genomic DNA from leaf samples

Young leaves from five accessions of *G. herbaceum* (A1-73, A1-79, A1-113, A1-Nisa, and A1-Wagad) and three of *G. arboreum* (A2-16, A2-26, and A2-113) were selected for analysis. Table 1 provides a detailed description of these accessions. For each accession, three independent replicates, with each replicate handled henceforth by a different researcher, were harvested resulting in a total of 24 individual samples.

**Table 1.** Description of the eight accessions used in the study.

| Species | Accession | Alternate Name | Sample Name | GRIN Accession<br>(PI) | SRA ID |
| --- | --- | --- | --- | --- | --- |
| *G. herbaceum* | A1-079 | VAR. KULJIANUM | A1-079 | PI 529655 | SRR8979915 |
| *G. herbaceum* | A1-Af | VAR. AFRICANUM | A1-Wagad | PI 630014 | SRR8979967 |
| *G. herbaceum* | A1-Nisa |  | A1-Nisa |  | SRR8979966 |
| *G. herbaceum* | A1-073 | M-881.8 | A1-073 | PI 485587 | SRR8979982 |
| *G. arboreum* | A2-016 | SMA-2 | A2-016 | PI 529719 | SRR8979962 |
| *G. arboreum* | A2-026 | KAPAS | A2-026 | PI 179607 | SRR8979976 |
| *G. arboreum* | A2-113 | SMA4 | A2-113 | PI 529740 | SRR8979927 |
| *G. herbaceum* * | A1-113 | LINE 4241 | A1-113 | PI 529686 | SRR8979908 |

\* incorrect placement suggests it is *G. arboreum* as per Grover (Grover *et al.*, 2021)

Nuclear genomic DNA was extracted using a combination of the nuclei isolation method of Paterson *et al*. (Paterson, Brubaker and Wendel, 1993) and the genomic DNA extraction method of the Qiagen Plant DNeasy Mini kit (Qiagen, Germantown MD). Briefly, for each of the 24 individual samples, a total of 200 mg of cotton leaf tissue was placed in liquid nitrogen in a mortar and ground into fine powder with a pestle. The tissue powder was suspended in 1.5 mL of ice-cold extraction buffer (Paterson, Brubaker and Wendel, 1993) and transferred into a microcentrifuge tube. The nuclei were pelleted by centrifugation at 4°C at 2,700x g for 20 min. After removing the supernatant, the pelleted nuclei were suspended in 400 μL of AP1 buffer (Qiagen Plant DNeasy Mini Kit) to which 4 μL of RNase A (100 mg/mL) was added (Qiagen, Germantown MD). The cell lysate was applied to a DNeasy Mini spin column. To get high molecular weight genomic DNA, the Qiagen column was centrifuged at 4,500x g for 1 min per the Qiagen Plant DNeasy Mini kit instructions. The plant nuclear genomic DNA was eluted from the column by adding 50 μL of 10 mM Tris-HCl buffer (pH 8.5). A NanoDrop One (ThermoFisher Scientific, Waltham MA) spectrophotometer was used to quantify the genomic DNA and determine 260/280 and 260/230 nm ratios. The DNA quality also was checked by running 100 ng of the genomic DNA sample on a 0.8% w/v agarose gel until the bromophenol blue marker dye reached about 55% of the gel length.

### 2.3 Preparation of the sequencing libraries

In ddRAD-Seq, genomic DNA is double digested using a pair of restriction enzymes; one of the enzymes is a rare cutter and the other is a frequent cutter. We ran egads on the genome sequence of *G. longicalyx* prior to digestion, and the results (Figure 2) suggested that a combination of the rare cutter *Hind*III and the frequent cutter *Sau*3AI (New England Biolabs, NEB, Ipswich, MA) would provide restriction fragments suitable for ddRAD-Seq (150-700 bp).

**Figure 2.**
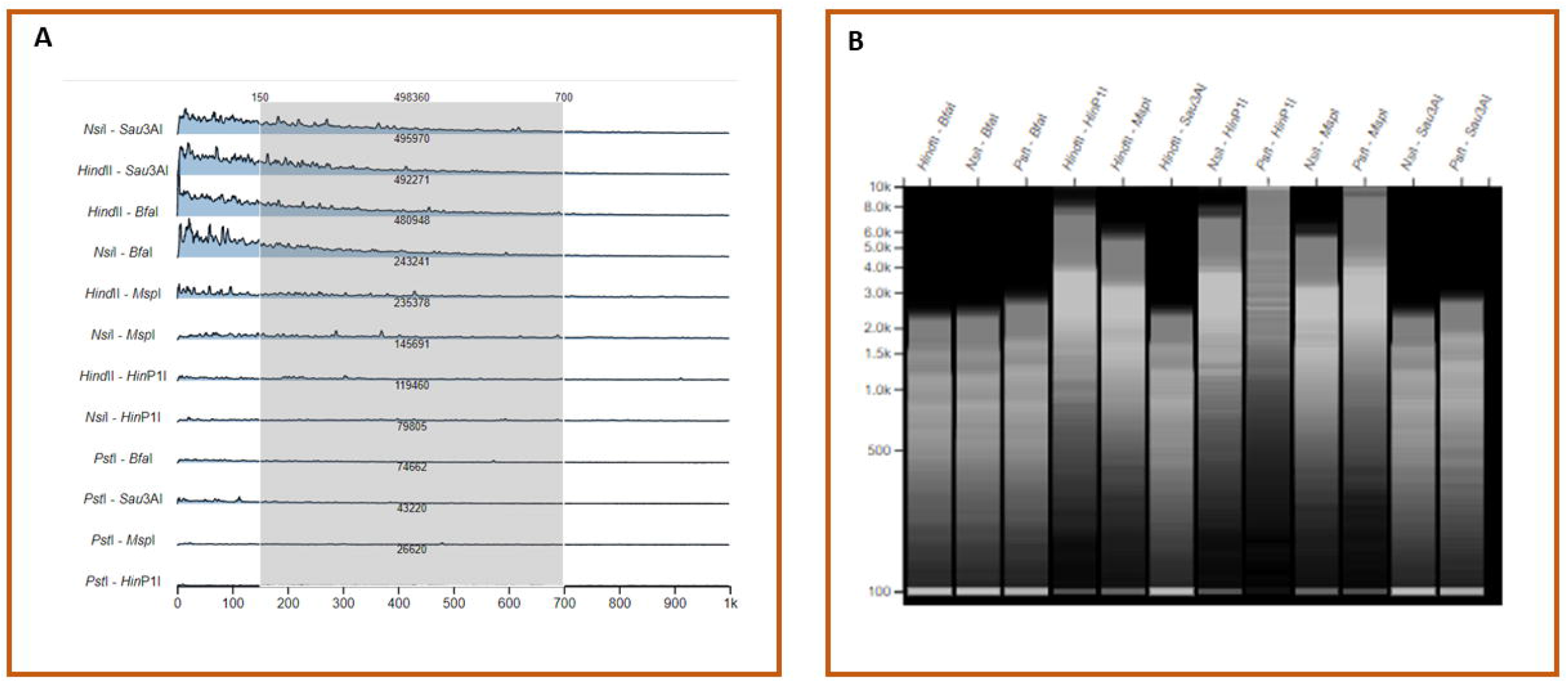
Outputs from egads. (A) Length distribution ridge plot of fragments between the enzyme pairs sorted by the number of fragments in selection area (highlighted in gray). (B) Gel image of double-digestion products as predicted and generated by egads.

For each individual sample, 450 ng of genomic DNA was digested for 2 hours at 37 ^0^C in 1X Cut Smart Buffer containing 0.8 units/μL *Hind*III and 0.2 units/μL *Sau*3AI in a reaction volume of 25 μL. Sixty nanograms of digested DNA was electrophoresed on a 1.5% w/v agarose gel to check for completeness of digestion. The restriction enzymes in a digested sample were not deactivated; rather, digestion products were immediately ligated or stored in the ligation-compatible Cut Smart Buffer at −20 ^0^C until ligation.

Overhang adapter pair sequences are modified Illumina adapter sequences that include a restriction enzyme overhang sequence. We purchased the following overhang adapter pair sequences from Integrated DNA Technologies (IDT, Coralville, IA):

*Hind*III adapters

5’-5Phos/AGCTTCTGTCTCTTATACACATCTGACGCTGCCGACGA-3’ and 5’-TCGTCGGCAGCGTCAGATGTGTATAAGAGACAGA-3’

*Sau*3A1 adapters

5’-5Phos/GATCCTGTCTCTTATACACATCTCCGAGCCCACGAGAC-3’ and 5’-GTCTCGTGGGCTCGGAGATGTGTATAAGAGACAG-3’

IDT synthesized and annealed the adapter pairs and added an HPLC purification step for increased efficiency. For each sample, 250 ng of *Hind*III-*Sau*3AI fragments were ligated with the *Hind*III overhang (2.5 pM) adapters and the *Sau3*AI overhang adapters (375 pM) in a 40 μL solution using 50 units of Quick Ligase (Quick Ligation Kit; NEB) that was 1X reaction buffer. Ligation was carried out at room temperature for 10 min.

The ligation products were purified immediately with AMPure XP beads (Beckman Coulter, Indianapolis, IN). The beads were equilibrated at room temperature for at least 30 min and vortexed for a few minutes before use. Thirty-two μL (0.8X volume of ligation reaction) of bead suspension was added to each ligation reaction, and the resulting mixture was vortexed briefly and then incubated at room temperature in a rotator for 5 min. The mixture was spun in a microcentrifuge at maximum speed for 1 min and allowed to further separate on a magnetic stand for several minutes. Once the supernatant appeared clear, it was carefully removed by pipette. While the tube remained on the magnetic stand, the pelleted beads were twice washed with 200 μL of freshly prepared 80% v/v ethanol. After the second wash, while retaining the tubes on the magnetic stand, all traces of ethanol were carefully removed. The tube cap was left open to let the pellet air-dry, but not to the point of cracking. To elute the ligated products, 15 μL of nuclease-free water was added to the tube. The tube was vortexed, gently mixed by rotation at room temperature for 5 min, spun in a microcentrifuge at maximum speed for 1 min, and placed on the magnetic stand until the supernatant was clear. The supernatant, which now contained the ligated products, was pipetted carefully into a new tube. One μL of this solution was quantified with the Qubit Fluorometer (ThermoFisher Scientific, Waltham, MA).

Polymerase chain reaction (PCR) was used to attach the i5 and i7 index primers [Nextera XT Index Kit v2 Set A (FC-131-2001) and Set B (FC-131-2002), Illumina, San Diego, CA] and to enrich for the ligated products. The index assignment for each sample is outlined in Supplementary File 1 (Tab 1). The reaction was composed of 0.12 μM each of the i5 and i7 primers, 30 ng of ligation products, 12.5 μL of Q5 High-Fidelity 2X Master Mix (NEB), and sufficient nuclease-free water to produce a final reaction volume of 25 μL. The reaction was initially denatured at 98 ^0^C for 30 seconds, which was followed by 14 cycles of denaturation at 98 ^0^C for 10 seconds, annealing at 65 ^0^C for 30 seconds, and extension at 72 ^0^C for 30 seconds. A final extension at 72 ^0^C for 5 minutes was performed to complete the polymerization of unfinished fragments.

Each of the 24 PCR reactions was cleaned with AMPure XP beads following the procedure stated earlier but with two modifications: (1) the bead suspension volume used was 25 μL (equal to the volume of the PCR reaction) and (2) the final product was eluted with 15 μL of EB buffer (10 mM Tris-HCl, pH 8.5). For each sequencing library, 1 μL was quantified with a Qubit Fluorometer while another 1 μL aliquot was run on the BioAnalyzer 2100 (Agilent Technologies, Santa Clara, CA) to check the library quality and determine the weighted average fragment size. The concentration from the Qubit reading (ng/ μL) and the weighted average fragment size (bp) were used to estimate the concentration of each library in nM using the formula: (concentration*10^6^)/(660*average fragment size). The first replicate from each of the eight individual plants was pooled in equimolar amounts to a final volume of 30 μL (R1), a second pool to generate the second replicate was similarly prepared (R2), and a third pool was likewise mixed to make the third replicate (R3). R1, R2 and R3 were size selected separately using SageELF (Sage Science, Beverly, MA) (Figure 1B). Fragments between 295 and 614 bp were retrieved from the selection step. Each of the three size-selected replicate pools was purified using 1.5X volume of AMPure XP beads following the modified protocol above. The Qubit and Bioanalyzer were again used to measure the concentration and weighted average of the fragments comprising each replicate pool after size selection. Equimolar amounts were pooled from each of the 3 replicates, and this was the input for Illumina sequencing.

### 2.4 Sequencing and analysis of sequencing reads

Libraries were sent to Novogene (Sacramento, CA) for sequencing on a single 2×150 lane of a HiSeq X Ten (Illumina, San Diego CA) and the raw sequencing reads were uploaded to the NCBI Sequence Short Read Archive (SRA) under the BioProject PRJNA690092 (Supplementary File 1; Tab 2). This ddRAD-seq dataset was combined with the dataset of whole-genome sequence (WGS) libraries of the same eight individuals (Table 1). The combined data-sets were mapped to the closely related outgroup species, *G. longicalyx* (Grover *et al*., 2020), using bwa v0.7.17-rgxh5dw (Li and Durbin, 2009) from Spack (Gamblin *et al*., 2015). Single-nucleotide polymorphisms (SNPs) were identified using the Sentieon software suite (Kendig *et al*., 2019) (Spack version sentieon-genomics/201808.01-opfuvzr), which is an optimization of the methods employed by the Genome Analysis ToolKit (McKenna *et al*., 2010). The DNAseq guidelines from Sentieon were followed, which include read deduplication, indel realignment, haplotyping, and joint genotyping. Parameters for mapping and SNP calling follow standard practices and are available in detail at https://github.com/IGBB/ddrad-methods/.

The multisample VCF was subsequently processed in vcftools (Spack version version 0.1.14-v5mvhea) (Danecek *et al*., 2011) to remove indels and to calculate the average read depth per sample. SNP sites were filtered to retain only those with <10% missing data, a maximum of two alleles, a minimum average read depth of 10, a maximum average read depth of 100, and a minor allele frequency of >5%. BCFtools was used to extract sample names, which were provided to vcftools to separate samples by library type (i.e., ddRADSeq or resequencing) and/or replicate. Phylogenetic trees were constructed using SNPhylo (Lee *et al*., 2014) and principal component analysis (PCA) was conducted using SNPRelate v 1.22.0 (Zheng *et al*., 2012) in R v4.0 (R Core Team, 2020). Data were visualized using ggplot2 (Wickham, Hadley (last), 2016).

## 3. Results and Discussion

### 3.1 Optimization of restriction enzyme pair for genomic DNA digestion

To take full advantage of the Illumina sequencing system, the restriction enzyme combination should produce DNA fragments between 150 and 700 bp because from our experience, fragments outside this range generate low quality sequence scores. To save time and resources, we developed egads (electronically guided digestion selection), available at https://github.com/IGBB/egads, a bioinformatics tool that predicts the likely size distribution of digestion products for a list of enzyme pairs for a genome sequence. Note that the sequence of the specific organism(s) being studied is not necessary for this to be useful; any related genome will provide candidate enzyme pairs, thereby reducing the number of combinations to test. Here we used egads on the *G. longicalyx* genome, which represents an outgroup to the sister species *G. herbaceum* and *G. arboreum*. The resulting output from egads is illustrated in Table 2 and Figure 2 (A and B), which suggests that combinations of *Nsi*I or *Hind*III (rare cutters) with *Bfa*I or *Sau*3A1 (frequent cutters) generate the greatest number of fragments in the desired range. Conversely, the results suggest avoiding any enzyme combination that includes *Pst*I due to the relatively low representation of *Pst*I sites in the *G. longicalyx* genome (1-2%). These results demonstrate the value of using egads to compare putative fragment size distributions generated by different enzyme pairs.

**Table 2.**
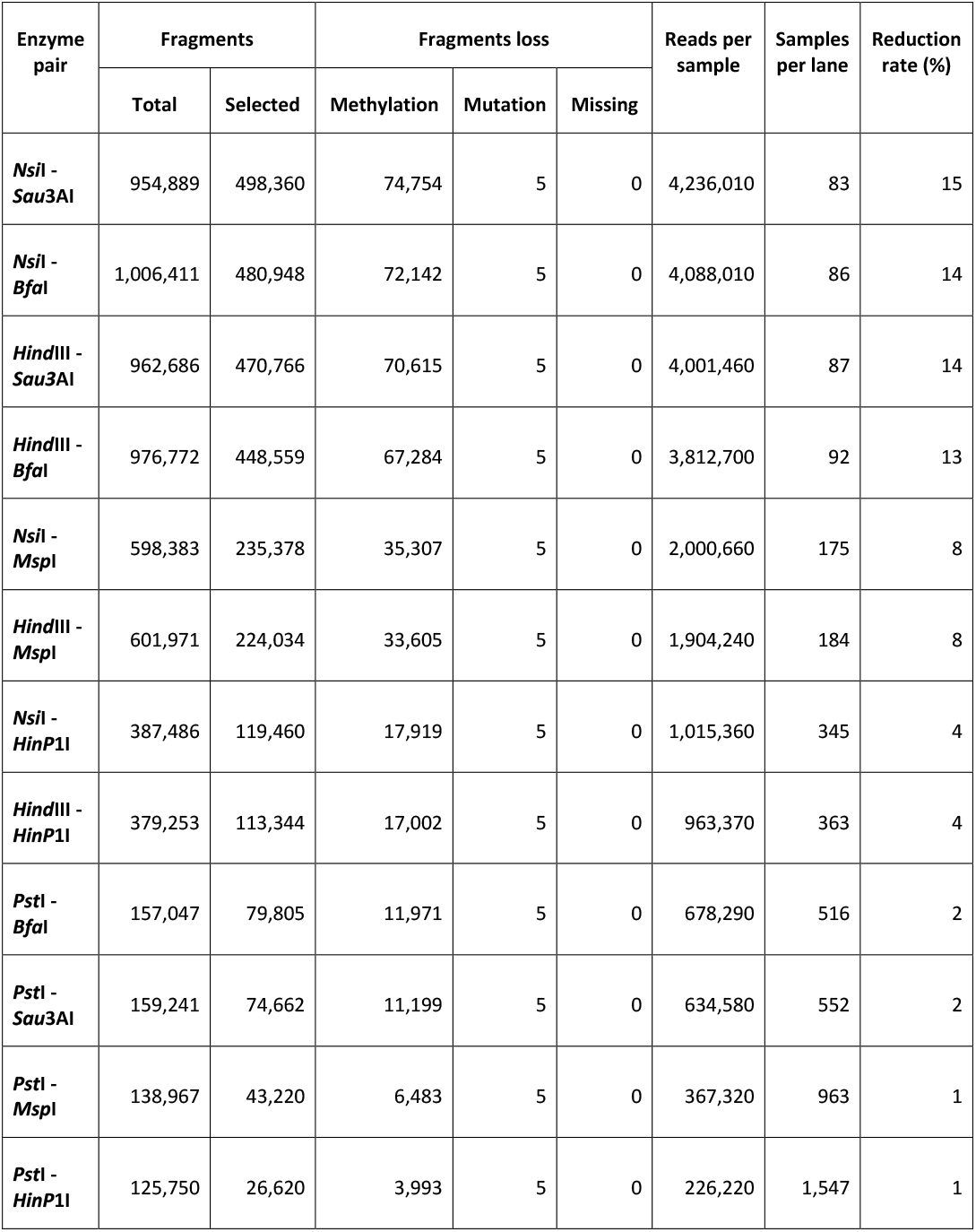
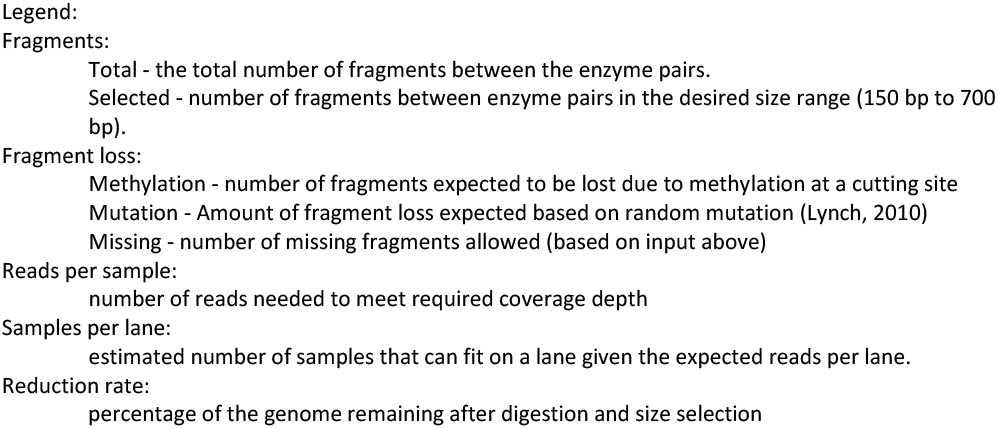
Enzyme pairs with output from egads and sorted by the number of selected fragments in the distribution plot.

| Enzyme pair | Fragments |  | Fragments loss |  |  | Reads per sample | Samples per lane | Reduction rate (%) |
| --- | --- | --- | --- | --- | --- | --- | --- | --- |
|  | Total | Selected | Methylation | Mutation | Missing |  |  |  |
| *NsiI* - *Sau3AI* | 954,889 | 498,360 | 74,754 | 5 | 0 | 4,236,010 | 83 | 15 |
| *NsiI* - *Bfal* | 1,006,411 | 480,948 | 72,142 | 5 | 0 | 4,088,010 | 86 | 14 |
| *HindIII* - *Sau3AI* | 962,686 | 470,766 | 70,615 | 5 | 0 | 4,001,460 | 87 | 14 |
| *HindIII* - *Bfal* | 976,772 | 448,559 | 67,284 | 5 | 0 | 3,812,700 | 92 | 13 |
| *NsiI* - *MspI* | 598,383 | 235,378 | 35,307 | 5 | 0 | 2,000,660 | 175 | 8 |
| *HindIII* - *MspI* | 601,971 | 224,034 | 33,605 | 5 | 0 | 1,904,240 | 184 | 8 |
| *NsiI* - *HinP1I* | 387,486 | 119,460 | 17,919 | 5 | 0 | 1,015,360 | 345 | 4 |
| *HindIII* - *HinP1I* | 379,253 | 113,344 | 17,002 | 5 | 0 | 963,370 | 363 | 4 |
| *PstI* - *Bfal* | 157,047 | 79,805 | 11,971 | 5 | 0 | 678,290 | 516 | 2 |
| *PstI* - *Sau3AI* | 159,241 | 74,662 | 11,199 | 5 | 0 | 634,580 | 552 | 2 |
| *PstI* - *MspI* | 138,967 | 43,220 | 6,483 | 5 | 0 | 367,320 | 963 | 1 |
| *PstI* - *HinP1I* | 125,750 | 26,620 | 3,993 | 5 | 0 | 226,220 | 1,547 | 1 |

Legend: Fragments: Total - the total number of fragments between the enzyme pairs. Selected - number of fragments between enzyme pairs in the desired size range (150 bp to 700 bp). Fragment loss: Methylation - number of fragments expected to be lost due to methylation at a cutting site Mutation - Amount of fragment loss expected based on random mutation (Lynch, 2010) Missing - number of missing fragments allowed (based on input above) Reads per sample: number of reads needed to meet required coverage depth Samples per lane: estimated number of samples that can fit on a lane given the expected reads per lane. Reduction rate: percentage of the genome remaining after digestion and size selection

### 3.2 Nuclear DNA quality, purity, and digestion

As with many genomics applications, ddRAD-Seq leverages high molecular weight DNA to generate reliable results. Several commercial kits are available for high molecular weight genomic DNA extraction, such as the Qiagen DNAeasy kit used here; however, direct use of these kits does not prohibit unwanted organellar DNA contamination. Therefore, to reduce contamination from cytoplasmic sources, buffered ground leaf slurries were enriched for nuclei prior to DNA extraction (see methods). All extractions generated high molecular weight genomic DNA with minimal or no smearing (Figure 3A), and the spectrophotometer absorbance ratios indicated high purity with minimal protein and ethanol (rinsing agent) presence. After DNA extraction, test digests were performed on a few samples using the four enzyme pair combinations identified by egads (data not shown). Based on the results, we chose *Hind*III (A’AGCTT) and *Sau*3AI (‘GATC) for ddRAD-seq because the pair (1) produces a suitable fragment size range, *i.e*., between 0.2 and 3 kb with a peak around 1 kb (Figure 3B), and (2) both exhibit optimal function under the same buffer conditions, which minimizes sample handling. Notably, these restriction enzymes also operate in a buffer that is neutral with Quick Ligase, further streamlining the protocol by eliminating the steps required to exchange the buffer for ligation. Therefore, careful consideration of the compatibility among enzymes and buffers, as well as their efficiency, can reduce the number of steps in the protocol and potentially cut significant time from the process, increasing overall efficiency.

**Figure 3.**
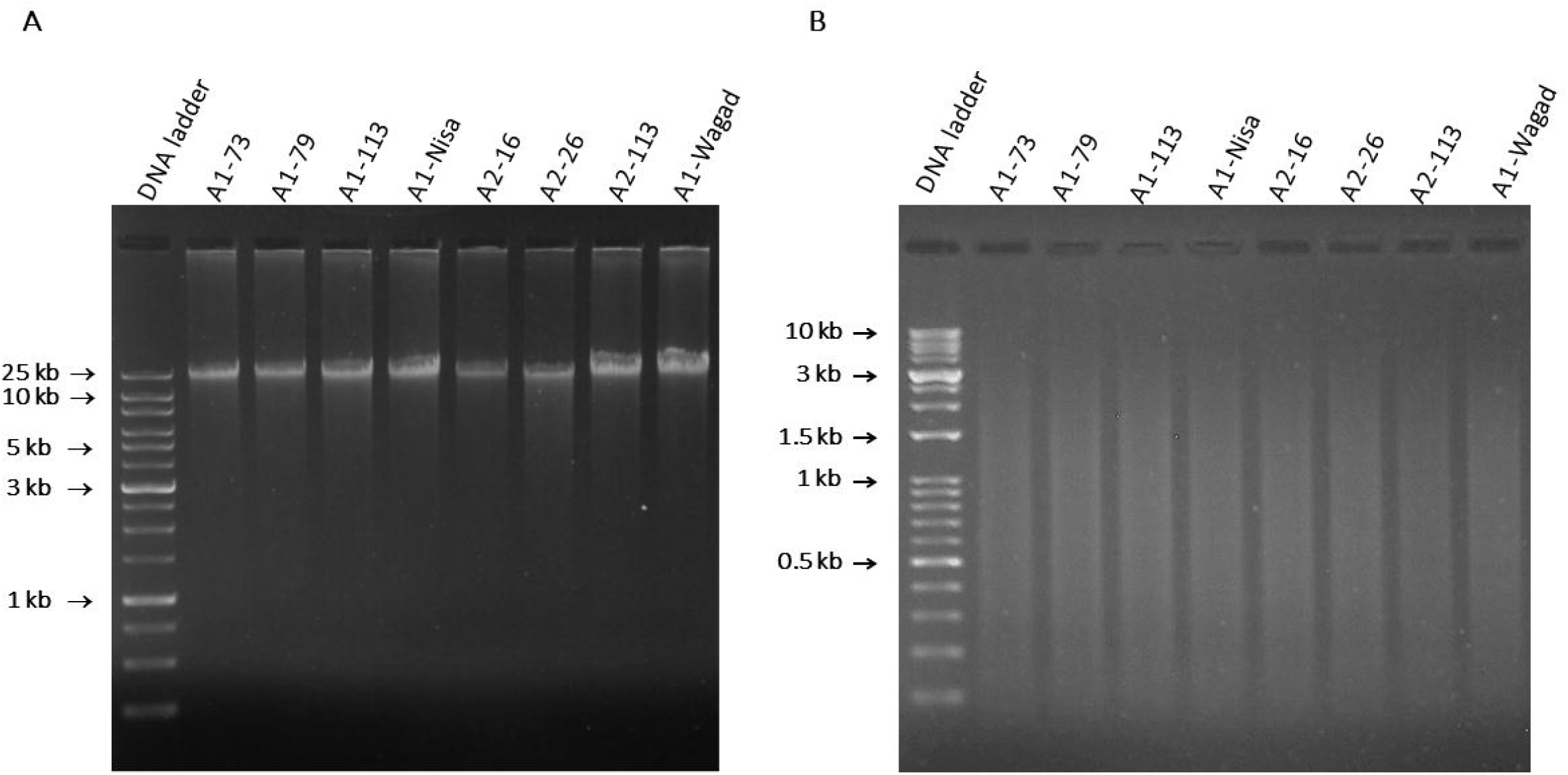
Agarose gel images of (A) extracted nuclear genomic DNA from ground leaves and (B) fragment profiles after double digestion with *Hind*III and *Sau*3A1.

### 3.3 Novel adapters and bead purification increase throughput and cost-effectiveness

Most of the previous work on ddRAD-Seq employs barcoded adapters, which was accomplished by synthesis of a unique adapter for each sample (or subset of samples) that often results in large adapter synthesis costs. Here, we have co-opted the sequences of the Illumina overhang adapters (forward and reverse) to function as the only two ddRAD-Seq adapters for all samples and employ complementarity to the commercially available Illumina-Nextera XT index primers to cost-effectively add barcodes to the adapter-ligated fragments via PCR. One of the main innovations here was the modification of the other end of these adapters (Illumina provided sequences) originally intended to generate 16S ribosomal RNA gene metagenomic libraries (see Methods) to instead be compatible with the cuts of the restriction enzymes selected. Here, the *Hind*III cut sequence (A’AGCTT) was added to the Illumina forward overhang adapter (in lieu of the 16S-compatible sequence), while the *Sau*3AI cut sequence (‘GATC) was attached to the Illumina reverse overhang adapter (see methods for full sequences). By using Quick Ligase, we were also able to cut the ligation time to 10 minutes at room temperature instead of overnight at 4 or 16 ^0^C. The Quick Ligase reaction chemistry was designed to have more collisions between the reactant molecules. This translates to decreasing not just the reaction time but also time spent by the researchers at the bench, resulting in better efficiency. Ligation of these overhang adapters then provides primer binding sites for the Illumina Nextera XT index primers, facilitating both library construction and indexing (Figure 1A).

Further optimizations were made by combining purification of the digested/ligated fragments with size selection using AMPure XP beads. Because the size range of DNA captured during bead purification is related to the concentration of the beads used (SPRIselect User Guide), bead purification can be used to exclude fragments outside of a desired size range while also minimizing DNA loss. Left side size selection, which theoretically eliminates fragments under 150 bp and simultaneously reduces the population of fragments of size 150 to 400 bp to preferentially retain fragments larger than 400 bp was performed using a bead to DNA volume ratio of 0.8X. This bead to DNA ratio was suitable to remove adapters, adapter dimers, very short fragments, and reactants in the samples tested here; however, we note that the target size range can be modified by altering the bead to DNA volume ratio. Larger fragments, for example, can be more stringently targeted by reducing the bead to DNA ratio because, as the relative concentration of beads decreases, the average size of the selected targets becomes larger and the size range narrower.

Another optimization used here combines library construction and barcoding in a single PCR-based step. We leveraged existing Illumina-Nextera XT index primers *(i.e*., i5 and i7 that are linked to sequencing primers and complementary to the adapter sequences) to differentiate among samples (Figure 1A). This further reduces both time and handling steps per sample by eliminating the cost and labor involved in synthesizing uniquely barcoded adapters for each sample or group of samples. Furthermore, because the i5 and i7 indices were tested by Illumina, the level of multiplexing is the same as in typical Illumina indexing (*i.e*., 384 unique combinations with dual barcoding), and the probability of barcode interaction leading to secondary structures is nearly absent. Also, because this protocol leads to individually barcoded samples (versus multi-level multiplexing; (Peterson *et al*., 2012), demultiplexing the sequences from pooled samples is straightforward.

A second round of bead cleanup purified the amplicons, this time using a 1:1 bead to DNA volume ratio to remove the primers and reagents while retaining amplicons between 200 and 600 bp. Interestingly, the size profiles of the replicates, which were produced independently by three researchers, were slightly different despite following the same protocol (Figure 4A), suggesting some variability in the bead handling steps. Although all were within the desired size range for Illumina sequencing, an optional secondary size selection for fragments between 295 and 614 bp (via SageELF) was included to further increase efficiency (Figure 1B). Figure 4B shows the Bioanalyzer output after the selection step and final bead purification steps (see methods).

**Figure 4.**
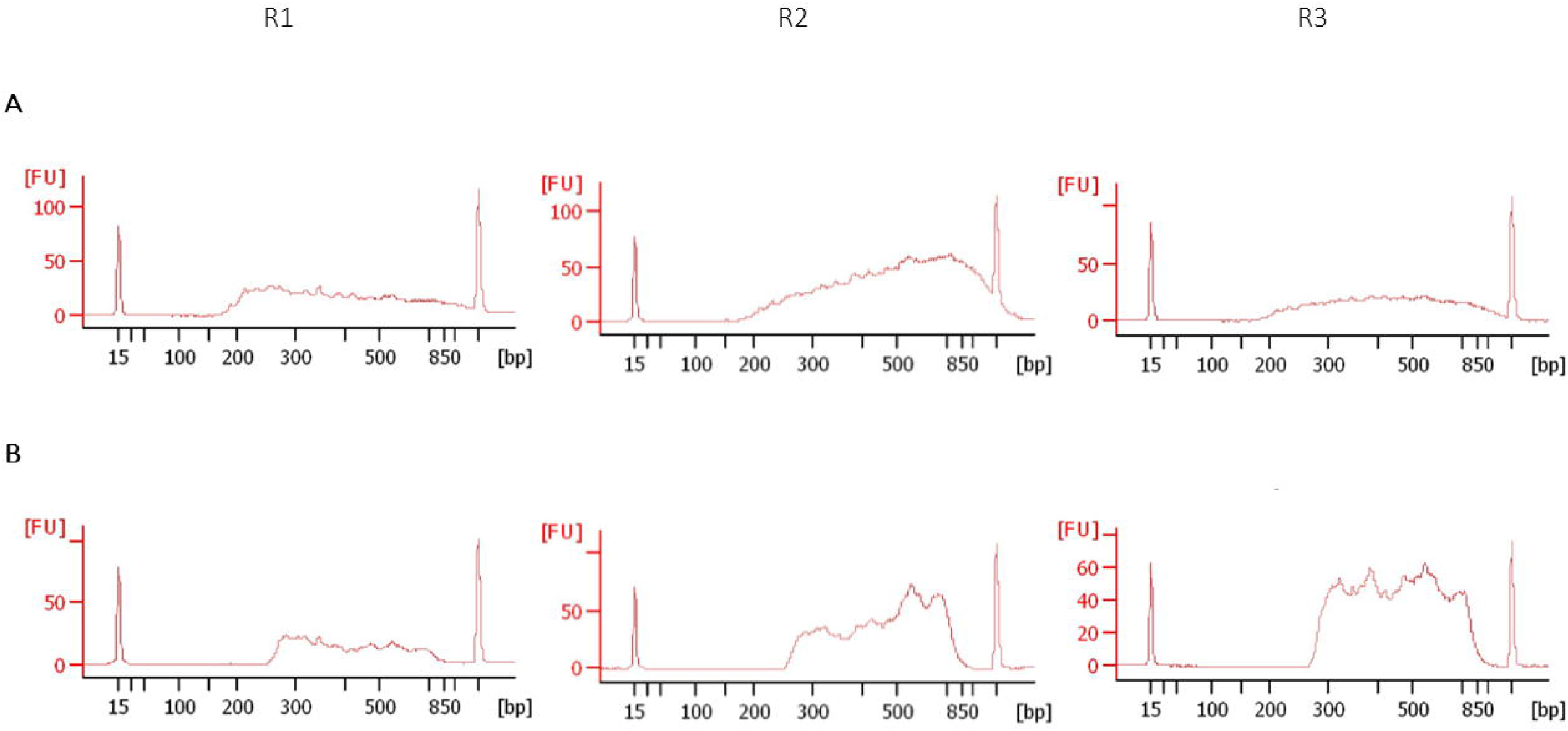
Bioanalyzer outputs (A) before and (B) after SageELF selection from three replicates (See Figure 1B).

### 3.4 SNP identification and consistency with genome resequencing

A major goal of ddRAD-Seq is to recover a common set of SNPs among samples useful for subsequent analyses (*e.g*., evolutionary analyses, phenotypic associations, etc.); however, it is important to note that accurate representation of the genome is critical. Here we evaluated the success of this protocol by characterizing the number and overlap of SNPs identified in our dataset relative to existing resequencing data for the same accessions. We selected five accessions of *G. herbaceum* (A1) and three accessions of *G. arboreum* (A2) which have existing resequencing data (Grover *et al*., 2021) to use in our ddRAD pipeline. A single lane of HiSeq X Ten produced 691.6 M reads from the 24 samples (8 accessions × 3 replicates), with an average of 28.8 M reads per sample. Given that the haploid genome size of both A1 and A2 is ∼1.7 Gb (Hendrix and Stewart, 2005), about 5% of the genome should be recovered by the restriction enzymes used, and a minimum of 2 million (M) reads would be necessary to cover the sequencing space by 10X or more at PE150. Accordingly, 691.6 M reads were sequenced from the 24 samples, with an average of 28.8 M reads per sample. Given this total output, about 173 individuals from this size genome could theoretically be genotyped to 20X coverage (4 M reads/sample). As expected, most of the reads fell within the projected length distribution acceptable for PE150 sequencing (Figure 5). Approximately 96% of reads per sample mapped to the reference (range: 19.6 M – 34.2 M, or 95 - 97% per sample; Figure 6), indicating good recovery. The actual genome reduction averaged 10.7% (range: 8.1% - 13.2%; filtering regions with three or fewer mapped reads) and drops to the expected reduction rate (∼5%) when filtering regions with less than 10 mapped reads. Of note, egads reports an expected reduction rate of 9% with size selection between 295 – 614 bp for the enzyme pair *Hind*III/*Sau*3AI.

**Figure 5.**
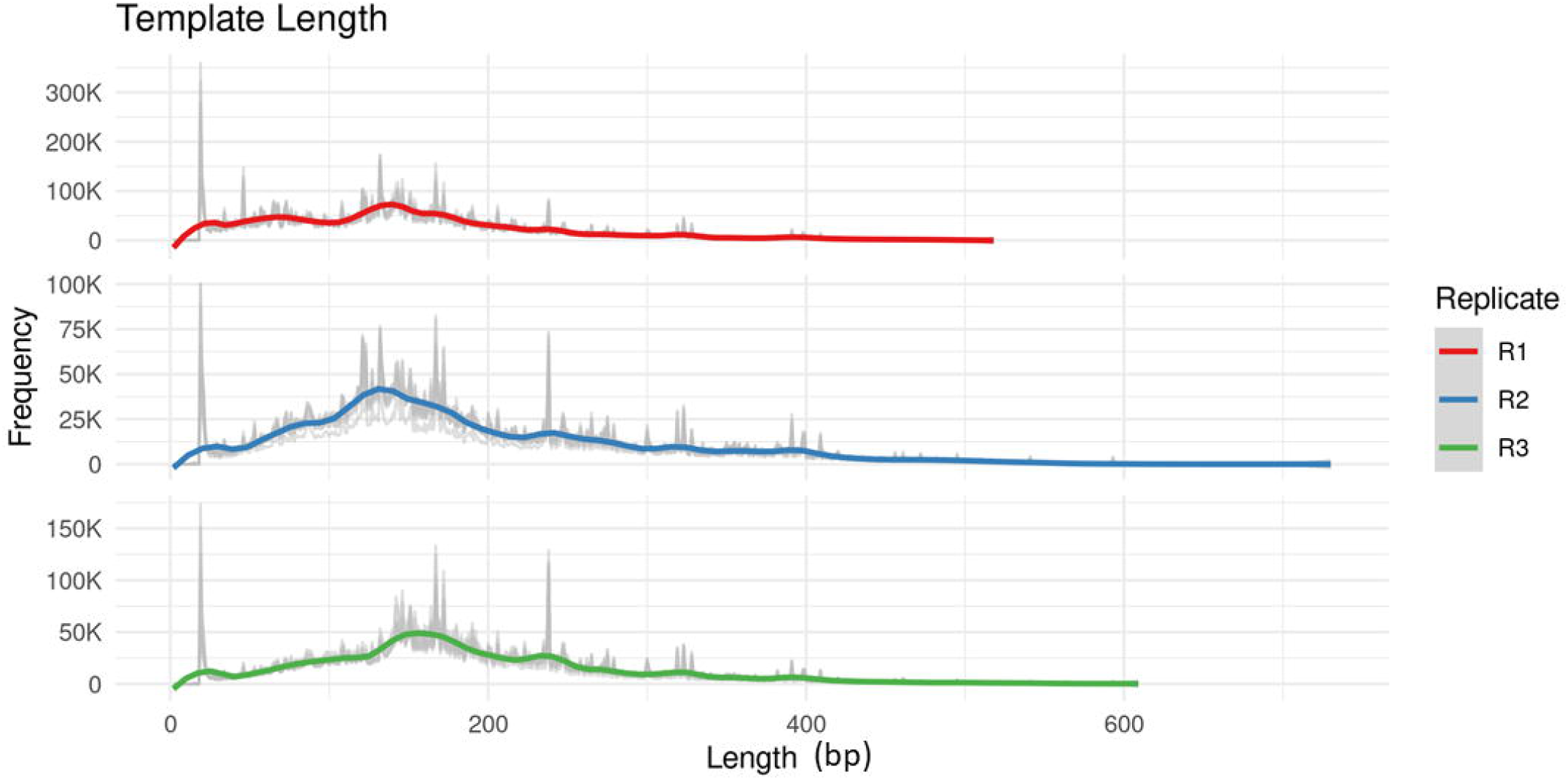
Distribution and frequency of the fragments of sequencing libraries arising from the three replicates after sequencing and quality control filtration from the reads.

**Figure 6.**
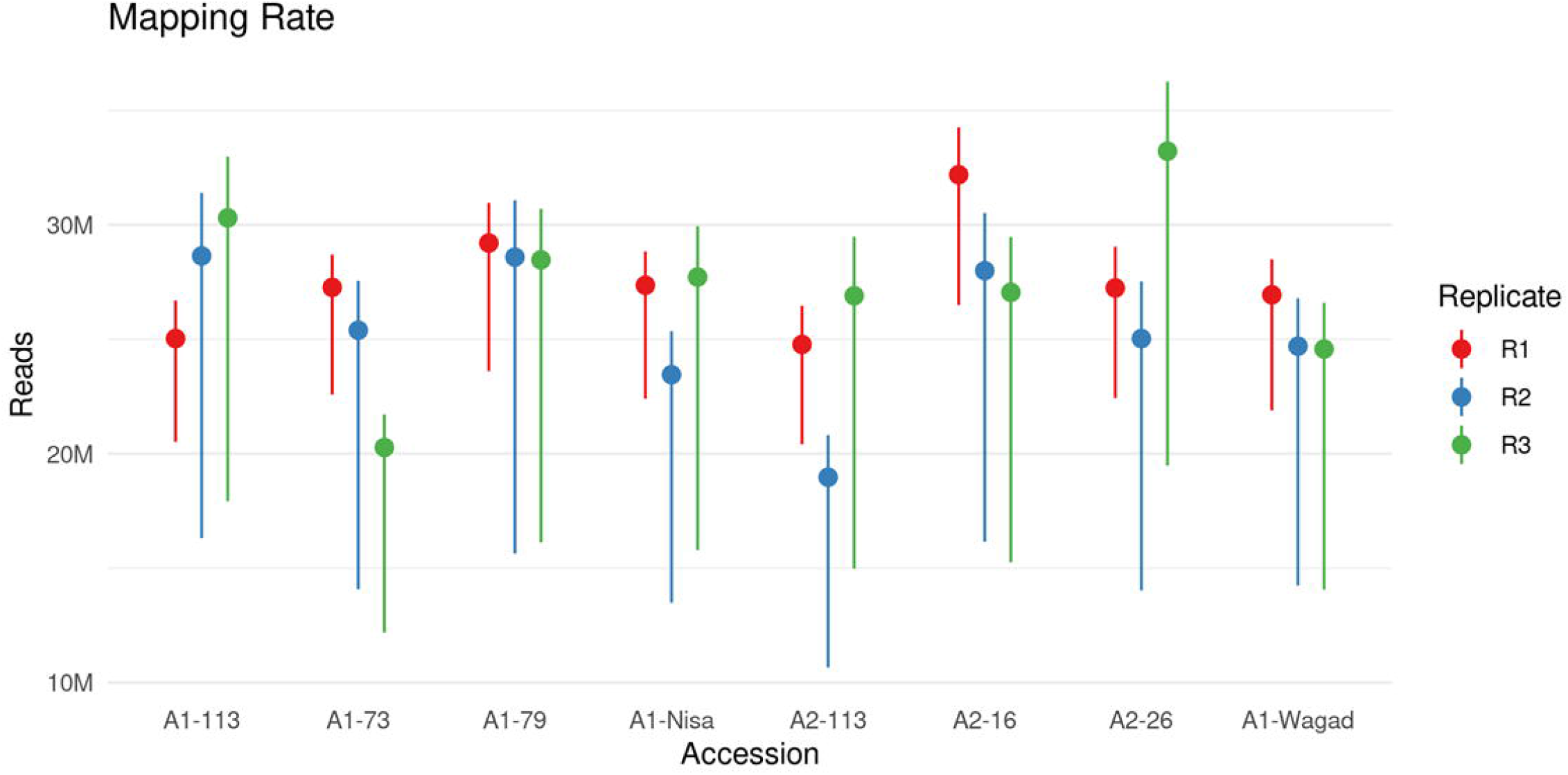
Graphic representation of the sequence reads that mapped into the reference *Gossypium longicalyx* genome. The top of each line represents the total number of reads, the point is the number of reads where both pairs aligned, and the bottom is the number of properly paired alignments, i.e., the read and its mate aligned to opposite strands of the same contig.

The ddRAD-seq data was assessed for potential bias by comparing it with high-coverage whole genome resequencing (WGS) for the same accessions (Grover *et al*., 2021). Joint-genotyping among samples (both ddRAD-Seq and WGS) identified 639,717 SNPs among the 8 resequenced and 24 ddRAD-Seq samples. Notably, our SNP detection rate is approximately 5.5% of the total SNPs previously detected by whole genome resequencing (∼12 million); (Huang *et al*., 2020; Grover *et al*., 2021) which parallels our expected reduction rate and suggests remarkable accuracy for our method. To further validate this method, the identified SNPs were subjected to principal component and phylogenetic analyses. Principal Component Analysis (PCA) of these data separates samples by species on principal component 1 (PC1), accounting for 7.22% of the variation among samples. Notably, A1-113 clustered with the other A2 samples, possibly indicating germplasm contamination and/or seed mislabeling, as previously noted for this accession (Grover *et al*., 2021). The second axis (PC2) separates the resequencing samples from the ddRAD-Seq replicates, accounting for 5.69% of the variation in the dataset (Figure 7A); however, phylogenetic analysis (Figure 7B) places all three replicates of each sample within the same clade as the genomic resequencing, indicating that they accurately represent the relationships among samples. These results indicate that the ddRAD-seq data, while capturing a slightly different representation of the SNPs contained within a sample, is consistent with the overall relationships among samples. Furthermore, the tight clustering of the ddRAD-Seq samples on the PCA suggests that this technique is highly replicable, even among different researchers.

**Figure 7.**
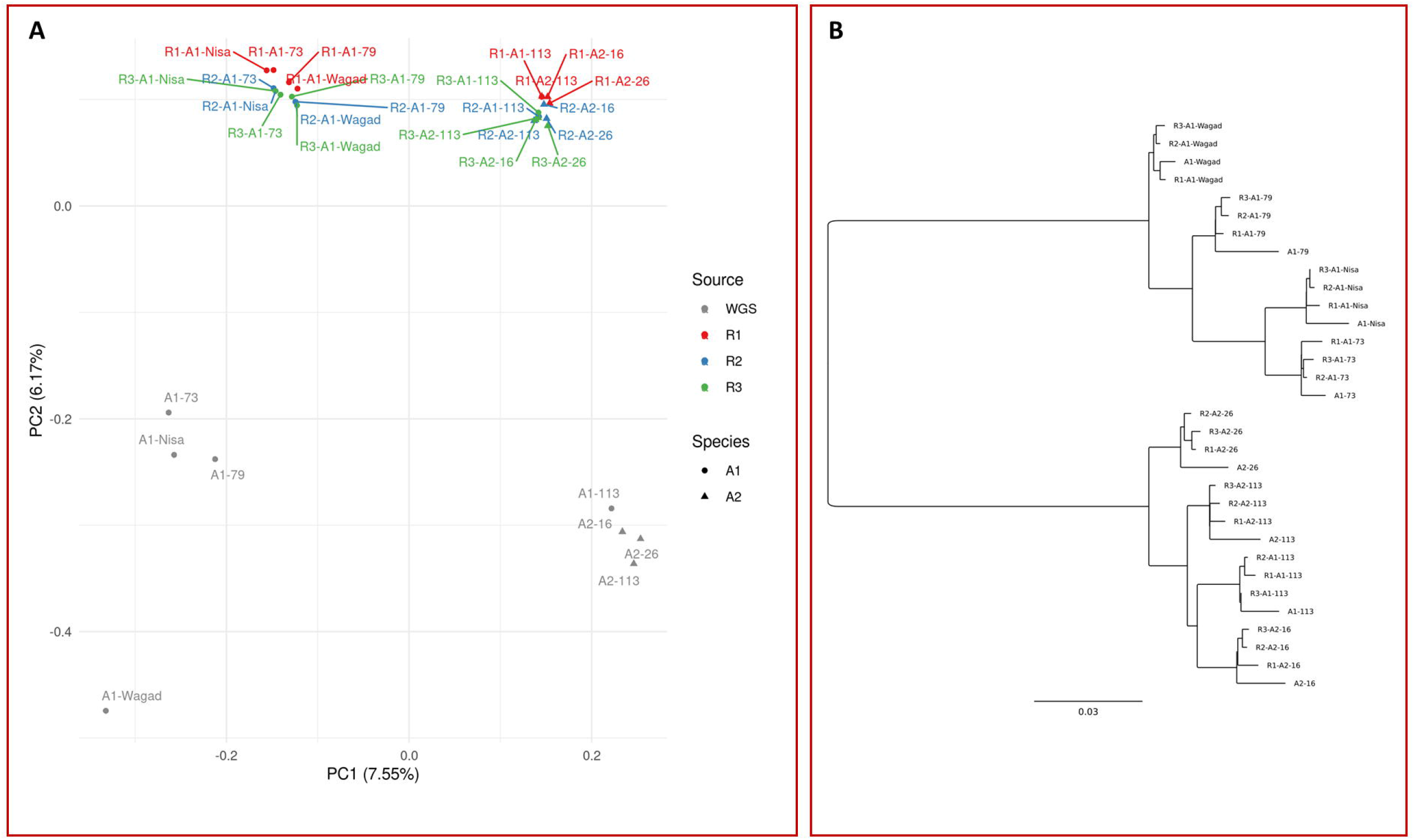
**A**. Principal component analysis and **B**. phylogenetic relationships of the SNPs derived from the resequenced genomes and ddRAD-Seq fragments.

We further explored the differences in SNPs detected by ddRAD-Seq by evaluating each SNP site using genomic resequencing as the expected outcome for that site (i.e., homozygous reference, homozygous derived, or heterozygous). Over 90% of expected homozygous sites (reference or derived) were accurately captured in the ddRAD-Seq data (Figure 8A), with only 1-2% of sites completely missed in the samples. Interestingly, the samples appeared to have “created” SNPs at 5 to 7% of the sites (Figure 8A), which is reflected in the “partial” and “incorrect” categories, suggesting novel SNPs that were detected in the heterozygous or homozygous states, respectively. While a small number of these may indicate somatic mutations captured in sequencing, most of these are likely attributable to the PCR amplification used to attach the indices and enrich for selected fragments, a process that is absent in generating PCR-free genomic resequencing libraries. These effects were also seen in comparing the expected heterozygous states among the samples, where just over 20% of heterozygous sites correctly identified both alleles in the ddRAD-seq and, in most cases, either the reference or alternate SNP was not recovered (Figure 8B). We note that PCR-based artifacts and/or biases (e.g., stochasticity, template switching, and polymerase errors) have been documented for multiple applications, including next generation sequencing (Acinas *et al*., 2005; Kebschull and Zador, 2015). Therefore, it is important to be cognizant of the potential for these errors and to consider methods for minimizing and/or accounting for them during analysis. For the present, while combining data sources arising from resequencing and ddRAD-Seq is not appropriate for research applications, consistency within the ddRAD-Seq samples and their reflection of the true relationships among the samples suggests that similarly prepared samples will be suitable for mutual analysis.

**Figure 8.**
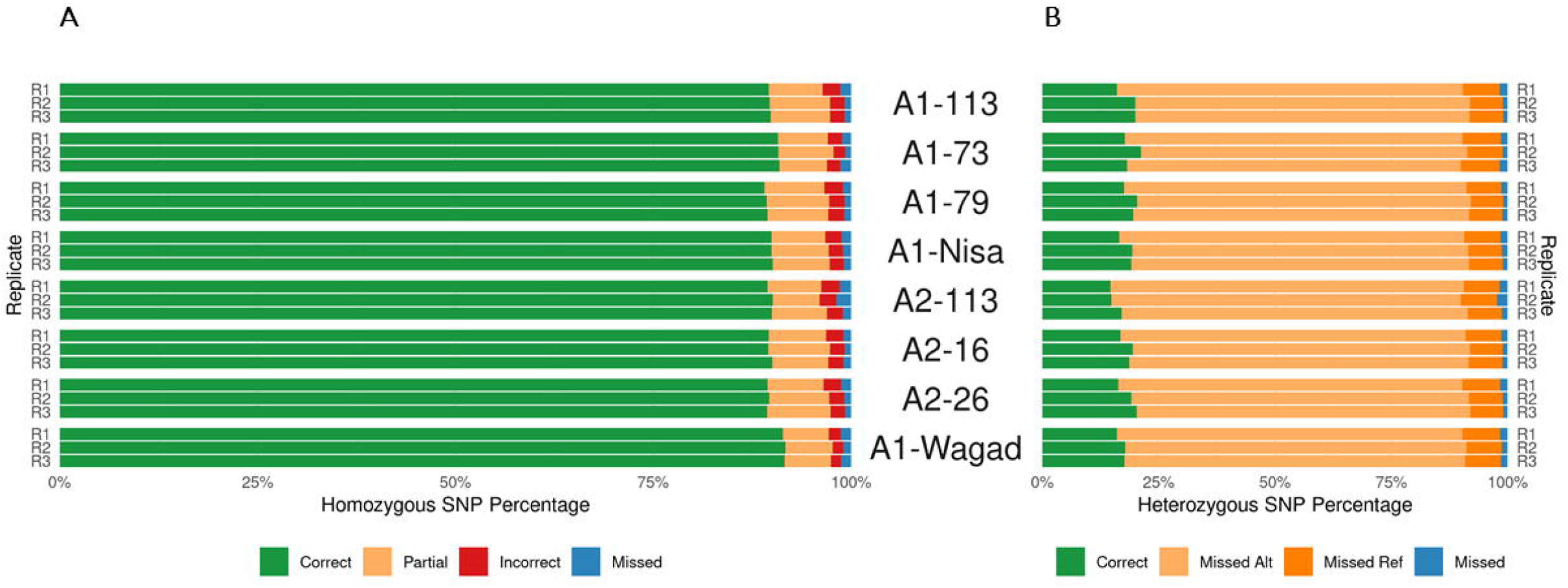
Categorization of ddRAD-Seq derived SNPs relative to whole genome resequencing. For each SNP site, the alleles predicted by the PCR-free, high coverage WGS was considered the expected state, and the proportion of ddRAD-Seq sites for that species/accession were categorized based how well they matched expectation. **A**. The proportion of correctly identified homozygous SNPs (ancestral or derived) is shown in green, and the proportion of sites that were completely absent in blue. Sites where novel alleles (i.e., presumed artifacts) were detected are pictured in orange (heterozygous, where one allele matched the expected state) and red (both alleles do not match the expected state). **B**. The proportion of sites where both heterozygous alleles were identified is shown in green, and the proportion of sites that were completely absent in blue. Sites where either the reference or alternate alleles were missed are shown in shades of orange.

## 4. Conclusions

Although next-generation sequencing costs continue to decline, the time and cost associated with this technology can still be prohibitive, particularly for population-based studies. Several methods have been developed to lower the cost and/or streamline next-generation sequencing for population-level studies; however, synthesizing unique barcodes for each sample or even a subgroup of samples is expensive. Moreover, multi-, or single-level multiplexing and demultiplexing of samples can be cumbersome, time consuming, and costly. Here we present an improvement of existing methods for ddRAD-Seq. Our ddRAD-Seq approach was successfully validated with whole genome resequencing and comparison to a high quality *de novo* genome sequence; however, our method should be applicable to any genome with or without a reference sequence. As we have illustrated, we used the genome sequence of *G. longicalyx* to predict the digestion products and to use as the reference for SNP detection in *G. herbaceum* and *G. arboreum*, sister taxa closely related to *G. longicalyx*. Our method (1) starts with high quality nuclear genomic DNA; (2) utilizes an algorithm that chooses the combination of a rare-cutting and a frequent-cutting restriction enzymes to digest the genomic DNA to the optimum sizes of the fragments for Illumina sequencing application; (3) employs reagents (e.g., Quick Ligase, compatible buffers) that significantly reduce reaction and processing times; (4) uses adapters that complement the Illumina-Nextera XT DNA i5 and i7 sets of indices, enabling the combination of barcode addition and library construction into one PCR step; and (5) takes advantage of magnetic bead to DNA stoichiometry to fine-tune the sizes of the selected fragments during clean up. Our comparison to whole genome resequencing data suggests that this method generates an accurate representation of the genome, resulting in SNPs suitable for downstream analyses (e.g., phylogenetics and association genetics) while simultaneously producing significant gains in cost and time.

## CRediT author statement

**ZVM:** Conceptualization, Methodology, Validation, Investigation, Resources, Writing-Original draft, - Review & Editing, Visualization. **CYH:** Conceptualization, Methodology, Validation, Investigation, Resources, Writing-Review & Editing, Visualization. **OP:** Validation, Investigation, Resources, Writing-Review & Editing. **MA:** Conceptualization, Software, Validation, Formal Analysis, Data Curation, Writing-Review & Editing, Visualization. **CEG:** Formal Analysis, Resources, Writing-Review & Editing. **DGP:** Conceptualization, Resources, Writing-Review & Editing, Funding Acquisition.

## Data availability

The data generated from this research is available as stated in Supplementary File 1.

## Acknowledgments

The authors thank the ResearchIT unit at Iowa State University for computational support.

